# The Alexander Disease Protein GFAP Drives Mitochondrial Fission

**DOI:** 10.1101/2024.02.20.581117

**Authors:** Ding Xiong, Linghai Kong, Ye Sing Tan, Fang Yuan, Zijun Sun, Xueyan Li, Emily Abella, Albee Messing, Su-Chun Zhang

## Abstract

Mitochondrial plasticity, coordinated by fission and fusion, is crucial to ensure cellular functions. Mitochondrial fission is mediated by the GTPase Drp1 at the constriction site, which is proposed to be driven by the actin-myosin contractile force. However, the mechanism that propels constriction remains unclear, and the potential involvement of additional mechanisms in this process remains an open question. Here, using structured illumination microscopy and electron microscopy, we show that the type-III intermediate filament glial fibrillary acidic protein (GFAP) closely surrounds mitochondria fission sites and associates with accumulated Drp1 molecules. Remarkably, loss of GFAP results in hyperfused mitochondria under physiological condition and even Ca^2+^-induced mitochondrial fission. Additionally, mutations of GFAP, the cause of Alexander disease, result in an elevated recruitment of Drp1 to GFAP, leading to significantly increased mitochondrial fissions. Taking together, these findings suggest a novel mechanism of mitochondrial division mediated by type-III intermediate filaments.

---

Mitochondria are highly dynamic in responding to cellular states, and this dynamic process is primarily regulated through fission and fusion^1^. Normal mitochondrial fission ensures cellular homeostasis, while excessive fission is often associated with major diseases, particularly neurodegenerative diseases^2,3,4^. Mitochondrial fission is a multi-step process in which the fission sites are marked by endoplasmic reticulum (ER) tubules followed by GTPase Drp1-mediated cleavage^5,6,7,8^. Given that the diameter of Drp1 spirals (30-50 nm) is significantly smaller than that of mitochondria (0.5-1 μm), a pre-constriction is necessary and actin filaments are proposed to facilitate constriction in a subset of fission routes ^9,10,11,12^. However, their necessity but not sufficiency with unclear mechanisms raises possible involvement of additional apparatuses.

Intermediate filaments (IFs) form elaborate networks, providing tensile strength and anchoring organelles, and demonstrate cell-type specific expression profiles^13,14,15,16,17^. In brain astrocytes, GFAP is the most abundant IFs. GFAP upregulation is nearly a universal feature under neurological conditions but the regulatory mechanism is unknown ^18^. Importantly, GFAP mutations result in astrocyte dysfunction, neuronal degeneration, and demyelination, leading to the often fatal Alexander disease (AxD)^19,20^. RNA sequencing of human astrocytes with AxD mutations revealed alterations in mitochondria function and metabolic pathway^21^, suggesting a role of GFAP in mitochondrial function. Although other IFs such as vimentin have been shown to interact with mitochondria^13^, whether and how IFs regulate mitochondria function remains unknown. Pathologically, how the disease-causing mutations in IFs affect mitochondria is an open question.

## Mitochondria align with GFAP filaments

To address these questions, we differentiated astrocytes from human pluripotent stem cells (hPSCs, H1 embryonic stem cell line) as previously described^22^. Following a 14-day maturation in the presence of bone morphogenetic protein 4 (BMP4) and ciliary neurotrophic factor (CNTF) (**Fig. 1A)**, the resulting astrocytes exhibited a star-shaped morphology with radial processes reminiscent of human astrocytes *in vivo* (**fig. S1A**). Using structured illumination microscopy (SIM), we observed GFAP filaments organized into a highly ordered fashion (**Fig. 1B**). These filaments exhibited a parallel alignment, forming thick bundles (0.52±0.13 μm in thickness under light microscopy, **fig. S1B**), and extended from the cell center to the processes **(Fig. 1B, box 1)**, while in the area between the astrocyte processes, fewer individual GFAP filaments showed a less directional orientation (0.18±0.02 μm) **(Fig. 1B, box 2)**.

**Figure 1.**
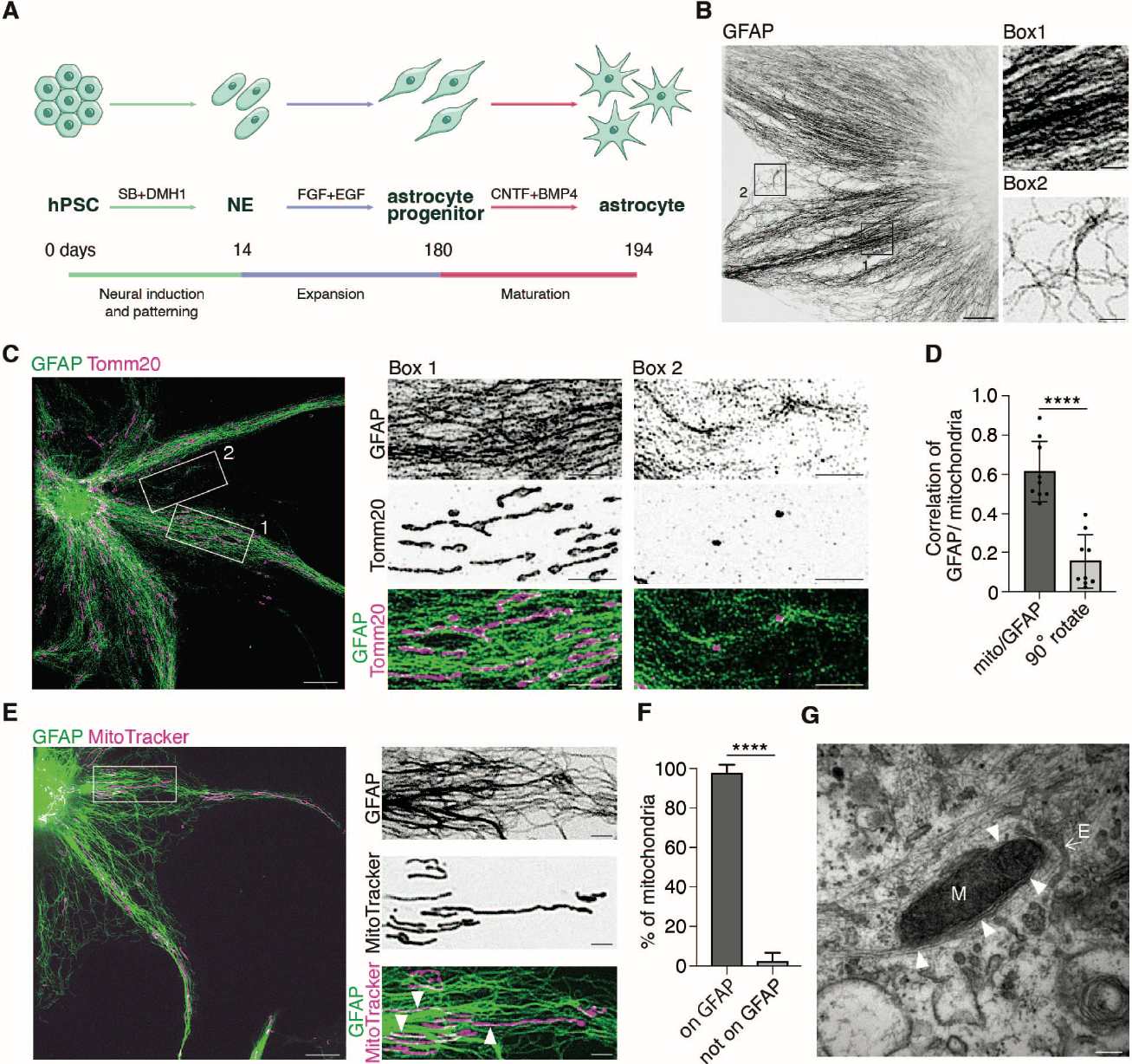
Spatial relationship between mitochondria and GFAP filaments. (**A**) Schematic of astrocyte generation from human PSCs (hPSCs). (**B**) Representative single-z SIM image of an astrocyte generated from H1 cell line immunostained with an antibody against GFAP. Boxes 1-2 mark regions of GFAP filaments at a higher magnification showing GFAP bundles (Box 1) and individual filaments (Box 2). (**C**) SIM image shows astrocyte co-immunostained with antibodies against GFAP (green) and Tomm20 (magenta). Area of GFAP bundles (Box 1) and individual filaments (Box 2) with Tomm20 are shown at higher magnification. (**D**) The level of colocalization between endogenous GFAP and mitochondria was determined by measuring the Mander’s coefficient for Tomm20 over GFAP, or when Tomm20 image was rotated 90° (****, p ≤0.0001, two-tailed paired *t*-test, n=9 cells). (**E**) Representative SIM image showing H1 astrocyte expressing GFAP-mNeon and stained with MitoTracker. An area of GFAP and mitochondria (white box) is shown in high-magnification images. Arrowheads indicate the positions of mitochondria aligned with GFAP filaments. (**F**) Quantification of the percentage of mitochondria aligned with GFAP filaments (196 mitochondria from 5 cells; p ≤0.0001, two-tailed paired *t*-test). (**G**) Representative transmission electron microscopy (TEM) image showing mitochondria (M), GFAP filaments (arrowhead) and ER (E, white arrow). Scale bar: SIM images, 10 μm; zoom-in SIM images, 2 μm; TEM image, 200 nm.

We next examined the spatial relationship between endogenous GFAP and mitochondria (labeled by Tomm20) and found that mitochondria were ‘embedded’ among GFAP filaments: they looked being wrapped by GFAP filaments and aligned in a highly parallel fashion along GFAP filaments, especially at the region with dense GFAP filaments (**Fig. 1C, box 1)**. This alignment was confirmed by the Mander’s coefficient quantification **(Fig. 1D)**. In regions with sparse individual GFAP filaments, there were few mitochondria. Nevertheless, they were still localized along GFAP filaments (**Fig. 1C, box 2**). We further confirmed such spatial distribution by visualizing GFAP via expression of WT-GFAP-mNeon under the GFAP promoter and mitochondria with MitoTracker (**Fig. 1E**). Quantitatively, majority of mitochondria aligned with the GFAP bundles, fewer with individual GFAP filaments, and only 2% of mitochondria appeared unassociated with GFAP (**Fig. 1F**). This colocalization was corroborated by transmission electron microscopy, through which we observed that mitochondria were wrapped by GFAP filaments (**Fig. 1G**). Thus, mitochondria closely align with GFAP filaments within human astrocytes.

### GFAP knockout inhibits mitochondrial fission

The close spatial vicinity between GFAP and mitochondria suggests that they may interact dynamically. Therefore, we performed live-imaging to assess mitochondrial dynamics in wildtype and GFAP knockout astrocytes. We generated a GFAP knockout cell line using the H1 embryonic stem cell by CRISPR, followed by differentiation into astrocytes **(fig. S2A-B)**. The absence of GFAP in the differentiated astrocytes was confirmed by western blot and immunostaining (**fig. S2C-D**). The astrocyte identity was validated by additional astrocyte markers, such as SOX9 and S100β (**fig. S2E**). Significantly, mitochondria underwent morphological change in GFAP knockout astrocytes. We categorized them based on size and shape characters as follows: fragmented (small or round), intermediate (medium size with round and tubular shape), and tubulated (long and higher interconnectivity)^23,24^. Among the isogenic control cells, the majority (79%) exhibited an intermediate mitochondrial phenotype **(Fig. 2A)**, with an average mitochondrial length of 10.3±2.7 μm **(Fig. 2C)**. Strikingly, in GFAP knockout astrocytes, the majority (83%) displayed tubular mitochondria with a significantly increased length at 37.1±8.1 μm, indicative of mitochondrial hyperfusion (**Fig. 2B-C)**.

**Figure 2.**
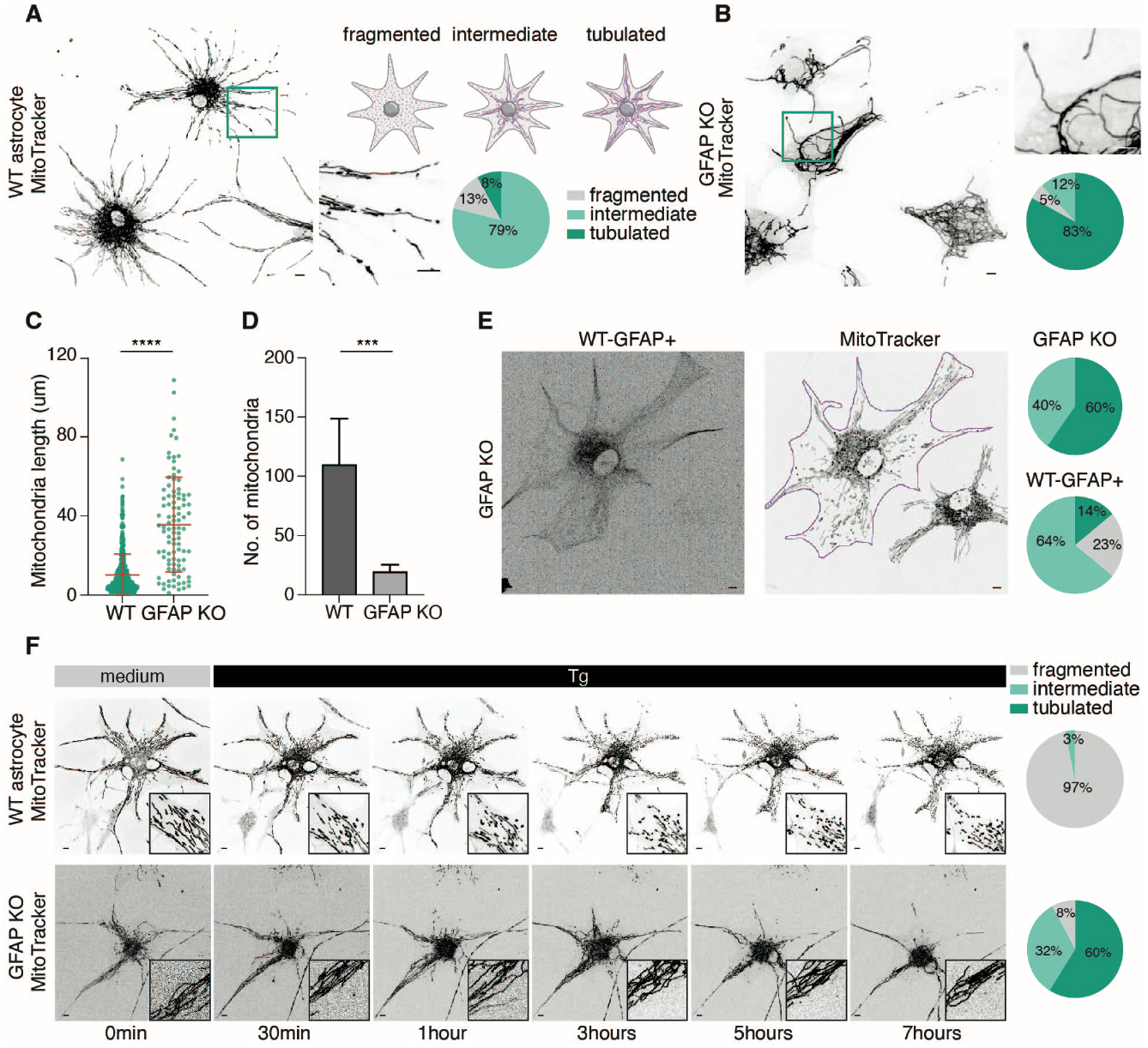
Inhibition of mitochondrial fission in GFAP knockout astrocytes. (**A**) Representative confocal image, magnified image of inset, and percentage of cells with indicated mitochondrial morphologies from wildtype astrocytes stained with MitoTracker (n=122 cells, 3 independent experiments). Mitochondria were visually scored as fragmented, intermediate or tubulated, as in the diagram. (**B**) Representative confocal image, magnified image of inset, and percentage of cells with indicated mitochondrial morphologies from GFAP knockout astrocytes stained with MitoTracker (n=144 cells, 5 independent experiments). (**C-D**) Quantification of individual mitochondrial length (**C**) and the number of mitochondria (**D**) in single astrocytes from WT and GFAP knockout astrocytes (549 and 97 mitochondria from 5 cells for WT and GFAP knockout cells, respectively). (**E**) Representative confocal images and percentage of cells with indicated mitochondrial morphologies in GFAP knockout astrocytes expressing WT-GFAP. Images showed GFAP KO astrocytes transfected with WT-GFAP-mNeon and stained with MitoTracker. Astrocyte with GFAP expression is outlined in purple. Percentage of cells with indicated mitochondrial morphologies from GFAP knockout and WT-GFAP expressing astrocytes from the same dish (149 GFAP KO cells and 83 GFAP expressing cells, 3 independent experiments). (**F**) Live-cell images and magnified images of WT and GFAP knockout astrocytes stained with MitoTracker when treated with 2 μM thapsigargin. Percentage of cells with indicated mitochondrial morphologies is shown at the right panel (62 WT and 119 GFAP KO cells, 3 independent experiments). ****, p ≤0.0001; ***, p≤0.001 by two-tailed unpaired *t*-test. Scale bar, 10 μm.

Correspondingly, the number of mitochondria in GFAP knockout astrocytes was drastically lower compared to that of control astrocytes **(Fig. 2D)**. The phenotype of mitochondrial hyperfusion in GFAP knockout astrocytes is likely due to the failure of fission or enhancement of fusion^6,25,26^, and indicates that GFAP critically regulates the balance of these antagonistic reactions. Indeed, expression of GFAP under a GFAP promoter rescued the phenotype in the GFAP knockout astrocytes, with a majority (64%) of astrocytes exhibiting an intermediate mitochondrial characteristic (**Fig. 2E**).

To discern whether the role of GFAP in mitochondrial dynamics is achieved through promoting fission or inhibiting fusion, we examined mitochondrial dynamics by an assay based on thapsigargin-stimulated mitochondrial fission. Thapsigargin triggers mitochondrial fission through the activation of calcineurin and subsequent dephosphorylation of Drp1 via Ca^2+^ release^27,28^. As a control, 97% of wildtype astrocytes underwent mitochondrial fission within 1-3 hours after treatment with thapsigargin **(Fig. 2F, movie. S1)**. If GFAP’s role is driving fission, thapsigargin treatment would not stimulate fission in the absence of GFAP; conversely, if GFAP functions by inhibiting fusion, thapsigargin treatment could stimulate mitochondria fission without GFAP. Interestingly, in the GFAP knockout astrocytes, mitochondria maintained the tubulated form and showed no response to thapsigargin stimulation for more than 7 hours (121/130 cells) **(Figure 2F, fig. S3A, movie. S1)**, indicating a loss of fission. Therefore, our data support that GFAP promotes mitochondrial fission instead of inhibiting fusion. To determine how loss of GFAP results in unsuccessful fission, we examined two possibilities: 1) the decrease in Ca^2+^ release and 2) loss of Drp1 accumulation. Live-imaging of cytosolic Ca^2+^ ruled out the possibility of Ca^2+^ release inhibition, as both GFAP knockout and control astrocytes showed acute Ca^2+^ increase immediately upon thapsigargin treatment and equilibrated to comparable level within 15min **(fig. S3B)**. However, immunostaining of Drp1 revealed decreased Drp1 puncta on mitochondria in GFAP knockout astrocytes **(fig. S3C)**, while no changes were observed in the fusion protein Mfn1/2 on mitochondria **(fig. S3D)**. These results suggest a potential role of GFAP in regulating mitochondrial fission machinery, likely through presenting Drp1 to mitochondria.

### Mitochondrial fission is enhanced by AxD mutations

In humans, GFAP mutations cause a rare yet often fatal Alexander disease (AxD), a complex leukodystrophy characterized by dysfunctional astrocytes which ultimately lead to neurodegeneration^19^. The above findings led us to hypothesize that mutant forms of GFAP disrupt mitochondrial distribution and fission/fusion dynamics in AxD patient astrocytes. We generated astrocytes from two AxD patients’ induced pluripotent stem cells (iPSCs) carrying the C88 and W416 mutations, as well as their respective isogenic control cell lines (R88 and R416) via CRISPR-mediated genetic correction^21^. In contrast to the radial distribution of mitochondria from the cell center to peripheral regions in isogenic control astrocytes, mitochondria in astrocytes with both C88 and W416 mutations exhibited a scattered distribution throughout the cytoplasm **(Fig. 3A)**. However, the mutation did not alter the alignment of mitochondria with GFAP filaments (**fig. S4**). Interestingly, in contrast to GFAP knockout astrocytes, the vast majority of AxD astrocytes (91%) displayed fragmented mitochondria, while the isogenic control astrocytes predominantly exhibited intermediate mitochondria (80% and 91% in R88 and R416 isogenic groups, respectively) **(Fig. 3A)**. Quantification of the length of individual mitochondria revealed notably shorter length in AxD astrocytes (3.9±1.1 μm for C88 mutant, 3.3±2.4 μm for W416 mutant) than in isogenic astrocytes (15.7±7.2 μm for R88 control, 10.3±2.2 μm for R416 control) **(Fig. 3B)**. Thus, GFAP mutations alter the distribution and fission/fusion dynamics of mitochondria.

**Figure 3.**
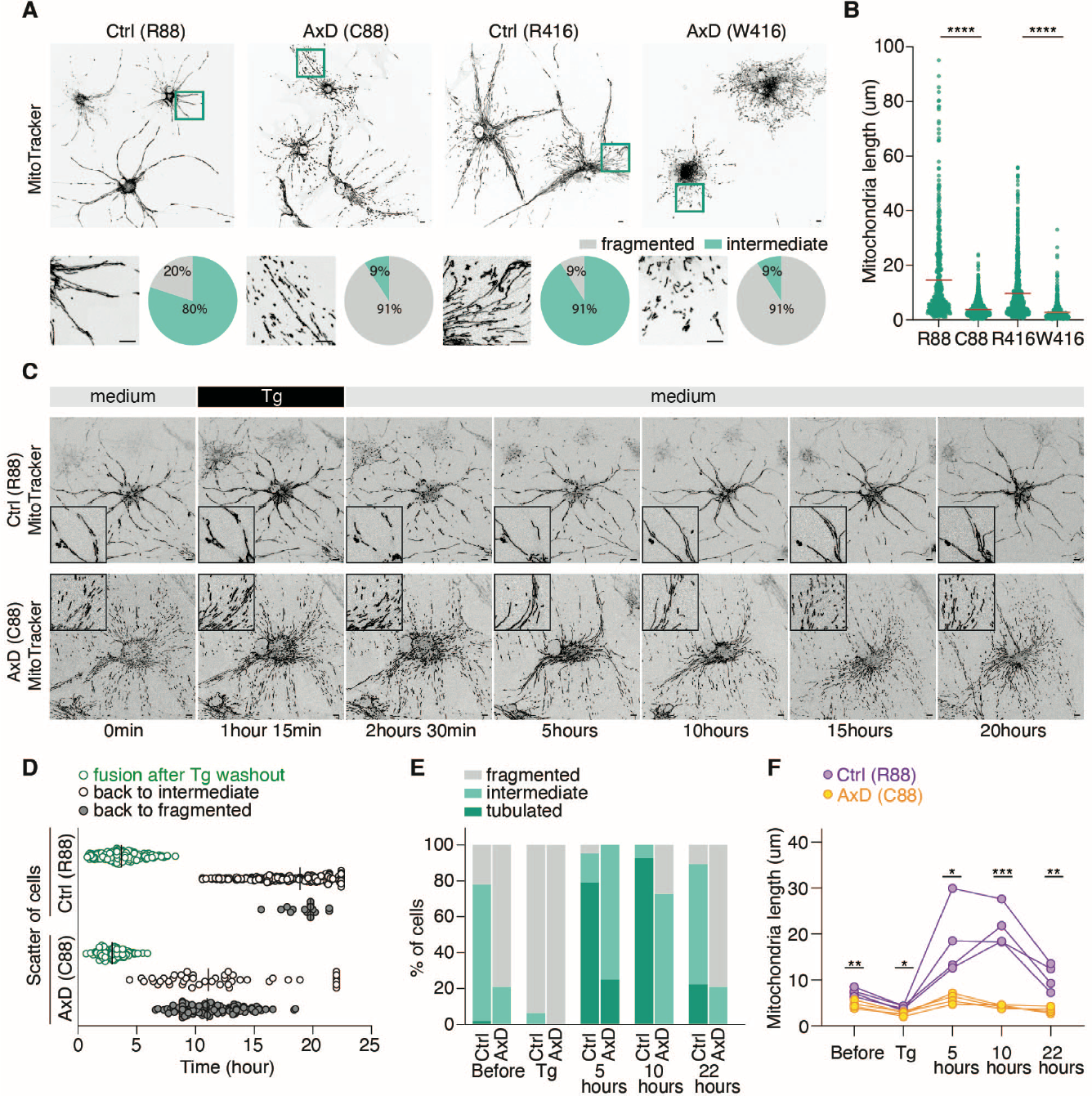
Mitochondrial fission is enhanced by AxD mutations. (**A**) Representative confocal images and percentage of cells with indicated mitochondrial morphologies in C88 and W416 AxD astrocytes and their isogenic controls R88 and R416. Astrocytes are stained with MitoTracker and insets are magnified below (n=188 R88 cells and 194 C88 cells from 11 independent experiments; n=140 R416 cells and 119 W416 cells, 4 independent experiments). (**B**) Quantification of individual mitochondrial length for both pairs of isogenic AxD astrocytes (n=524, 1705, 770 and 650 mitochondria from 5 cells for R88, C88, R416 and W416, respectively). (**C**) Live-cell images showing the dynamics of mitochondrial fission-fusion in control (R88) and AxD (C88) astrocytes. Astrocytes stained with MitoTraker are synchronized by 2 μM thapsigargin and exchanged back to normal medium as indicated. (n=186 R88 cells and 151 C88 cells, 3 independent experiments). (**D**) Scatter plot shows the time point at which mitochondria start fusion after thapsigargin washout (green), return back to intermediate mitochondria status (black circle) or further go back to fragmented status (grey). Each dot represents one cell, with average time shown as black lines. (**E**) Percentage of R88 and C88 astrocytes in tubulated (green), intermediate (light green) and fragmented (grey) status at indicated time. 92% (172/186) control astrocytes and 25% (37/151) AxD astrocytes form hyperfused mitochondria before return back to intermediate or fragmented status. (**F**) Quantification of individual mitochondrial length for R88 and C88 astrocytes at indicated time. One dot means averaged mitochondria length from one cell (4 cells for R88 and C88 cells, respectively). The plot for the length of individual mitochondria is shown in fig. S5B. *, p<0.05, **, p<0.005, ***, p<0.001 by two-tailed unpaired *t*-test. Scale bar, 10 μm.

Next, we investigate how mitochondrial dynamics is altered by the mutant GFAP. We synchronized the mitochondrial state by thapsigargin treatment followed by its washout. We found no difference between AxD and isogenic control cells in thapsigargin induced mitochondrial fragmentation (90/94 cells in R88 group, 146/152 cells in C88 group; 91 /101 in R416 group, 85/97 in W416 group) **(fig. S5a, movie. S2)**, indicating that the fission machinery is functional in both AxD and isogenic control astrocytes. After thapsigargin withdrawal, mitochondria elongation started in ∼3 hours in both AxD and control astrocytes (3.8±1.6 hours in R88 group, 2.9±1.0 hours in C88 group) **(Fig. 3C-D)**, suggesting that the mitochondrial fusion machinery is unlikely affected by the GFAP mutation. Over a 22-hour tracking period, 92% control astrocytes formed hyperfused mitochondria (in agreement with previous study in rat liver cells^27^), whereas only 25% AxD astrocytes formed hyperfused mitochondria and the remaining fused to “intermediate” mitochondria **(Fig. 3C, movie. S3)**. Besides, after mitochondrial elongation, the majority of AxD astrocytes rapidly reverted to fragmented mitochondria at ∼12 hours, much earlier than the control astrocytes which transitioned back to the intermediate state at ∼18 hours **(Fig. 3D)**. These dynamic changes were also reflected by the length of individual mitochondria within single cells analyzed at five time points, indicating that GFAP mutation correlates with an enhanced mitochondrial fission activity **(Fig. 3E-F, fig. S5B)**. A similar phenotype was also seen in astrocytes with the W416 mutation **(fig. S5C-E)**, indicating that the increased fission is a consistent phenotype from AxD GFAP mutations. Thus, GFAP mutations primarily upregulate mitochondrial fission.

### GFAP filaments directly participate in mitochondrial constriction

To uncover the mechanisms underlying the role of GFAP in mitochondrial fission, we assessed their spatiotemporal interaction using structured illumination microscopy. we noticed that mitochondria with a narrowing diameter is at the site where GFAP filament crossed (**Fig. 4A)**. This observation was corroborated by electron microscopy, revealing the intertwining of the ER tubules and GFAP filaments around the constricted “neck” of dividing mitochondria (**Fig. 4B**), which suggests that GFAP filaments may participate in mitochondrial constriction. By live-cell imaging of astrocytes expressing the fluorescent-tagged GFAP (WT-GFAP-mNeon or mutant C88-GFAP-mNeon) (**fig. S6A)**, we observed that GFAP filaments crossed and scanned/slid along mitochondria during the constriction process (**Fig. 4C-G, fig. S6B-D, movie. S4-5**). Notably, the constriction of mitochondria was spatiotemporally coupled with the movement of GFAP filaments along mitochondria (**Fig. 4D-E, fig. S6B, movie. S4**). Further quantification showed that majority of fission events (83% in WT-GFAP expressing astrocytes and 86% in C88-GFAP expressing astrocytes) occurred at locations where GFAP filaments intersected with mitochondria. This was confirmed by measuring the fluorescence intensity of GFAP and mitochondria perpendicular to the fission sites (**Fig. 4D, G**). The tight association of GFAP filament with mitochondria covering the constriction process indicates that intermediate filament-mitochondria contact is important particularly at the early, constriction stage. In contrast, when mitochondria were aligned in parallel with GFAP, we observed few fission events (**fig. S6E**). These results indicate that GFAP filaments participate in mitochondrial constriction during the fission process.

**Figure 4.**
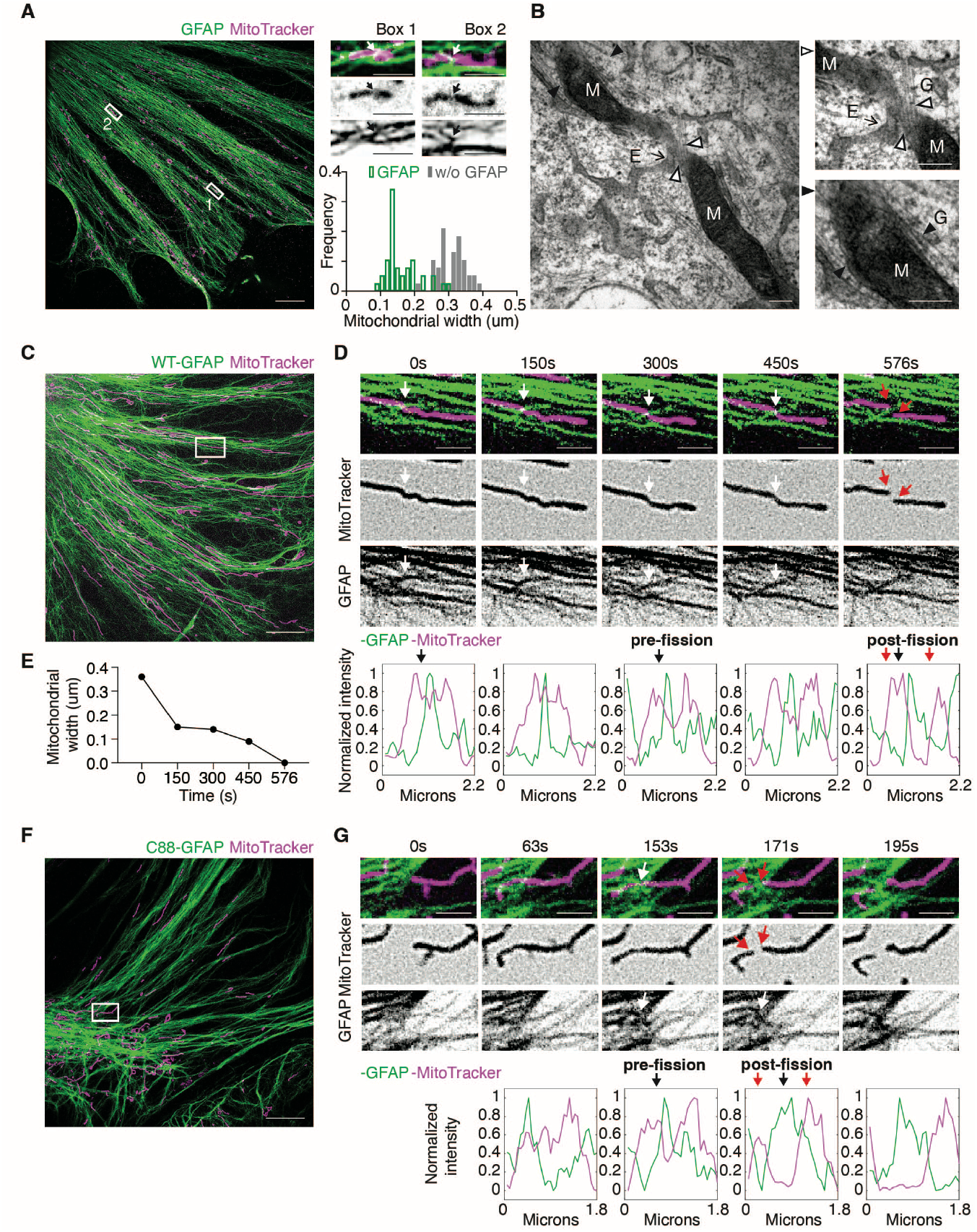
GFAP filaments directly participate in mitochondrial constriction. (**A**) Representative SIM image and histogram for mitochondrial diameter of R88 control astrocytes immunostained for GFAP (Cell signaling technology, 34001) and stained with MitoTracker. Insets are magnified on the top right. Arrows indicate the regions where GFAP filaments cross mitochondria. Histogram at the bottom right showing diameter of mitochondria with or without GFAP crossing (with GFAP: n=38 events, without GFAP: n=38 events, 8 independent experiments) (**B**) Representative transmission electron microscopy (TEM) image with zoom-in panels showing GFAP filaments aligned in parallel (black arrowhead) with mitochondria (M) or crossing (white arrowhead) mitochondria next to ER (E). (**C**) SIM image of a live astrocyte expressing WT-GFAP-mNeon and stained with MitoTracker. (**D**) Upper, high-magnification images of the selected region in (C) showing GFAP filament crossing the site of mitochondrial division (white arrow) before fission (red arrow). Dashed line indicates the mitochondrial region used to generate line scan plot. Bottom, corresponding line scans analyze the fluorescence intensity of mitochondria and GFAP pre- and post-fission. Black and red arrows on line scan analysis correspond to white arrow and red arrow positions at pre-constriction and division sites, respectively (44/53 mitochondria with WT-GFAP crossing, 4 independent experiments). (**E**) Quantification of corresponding mitochondrial width at the position of GFAP crossing (white arrows) in (D). (**F-G**) As in (C-D), SIM images, high-magnification and line scan analysis of a live astrocyte expressing C88-GFAP-mNeon and stained with MitoTracker (24/28 mitochondria with C88-GFAP crossing, 3 independent experiments). Scale bar: SIM images, 10 μm; zoom-in SIM images, 2 μm; TEM image, 200 nm.

### GFAP filaments serve as scaffolds for fission protein Drp1

Mitochondrial constriction is followed by Drp1 GTPase-mediated cleavage to achieve fission^5^. The fact that both wildtype and mutant GFAP filaments exert constriction similarly (**Fig. 4**) but AxD patient astrocytes exhibit more pronounced mitochondrial fission (**Fig. 3**) raised a question of whether GFAP interacts with fission molecules and GFAP mutations alter such interaction. Under structured illumination microscopy, we found that Drp1 predominantly localized in close proximity or tethered to GFAP filaments (**Fig. 5A**), suggesting that GFAP filaments may act as an anchor for Drp1 oligomerization and promote the local enrichment of Drp1 which drives the fission process. Quantitative analysis revealed an increased number of Drp1 puncta localized to mutant GFAP compared to wildtype GFAP (**Fig. 5A-B**). Immunoblotting confirmed that the increased binding in AxD astrocytes is not due to an increase in the total Drp1 level (**Fig. 5C**). Furthermore, at the mitochondrial constriction sites, Drp1 puncta are highly co-localized with the regions where GFAP filaments intersected mitochondria (**Fig. 5D**), while a remarkable reduction of Drp1 puncta was observed on mitochondria in GFAP knockout astrocytes (**fig. S3C**). Taking together, these data suggest that GFAP filaments recruit or present Drp1 to mitochondria, and this hypothesis also explains the phenotype of more mitochondrial fission events observed in AxD astrocytes **(Fig. 3)**.

**Figure 5.**
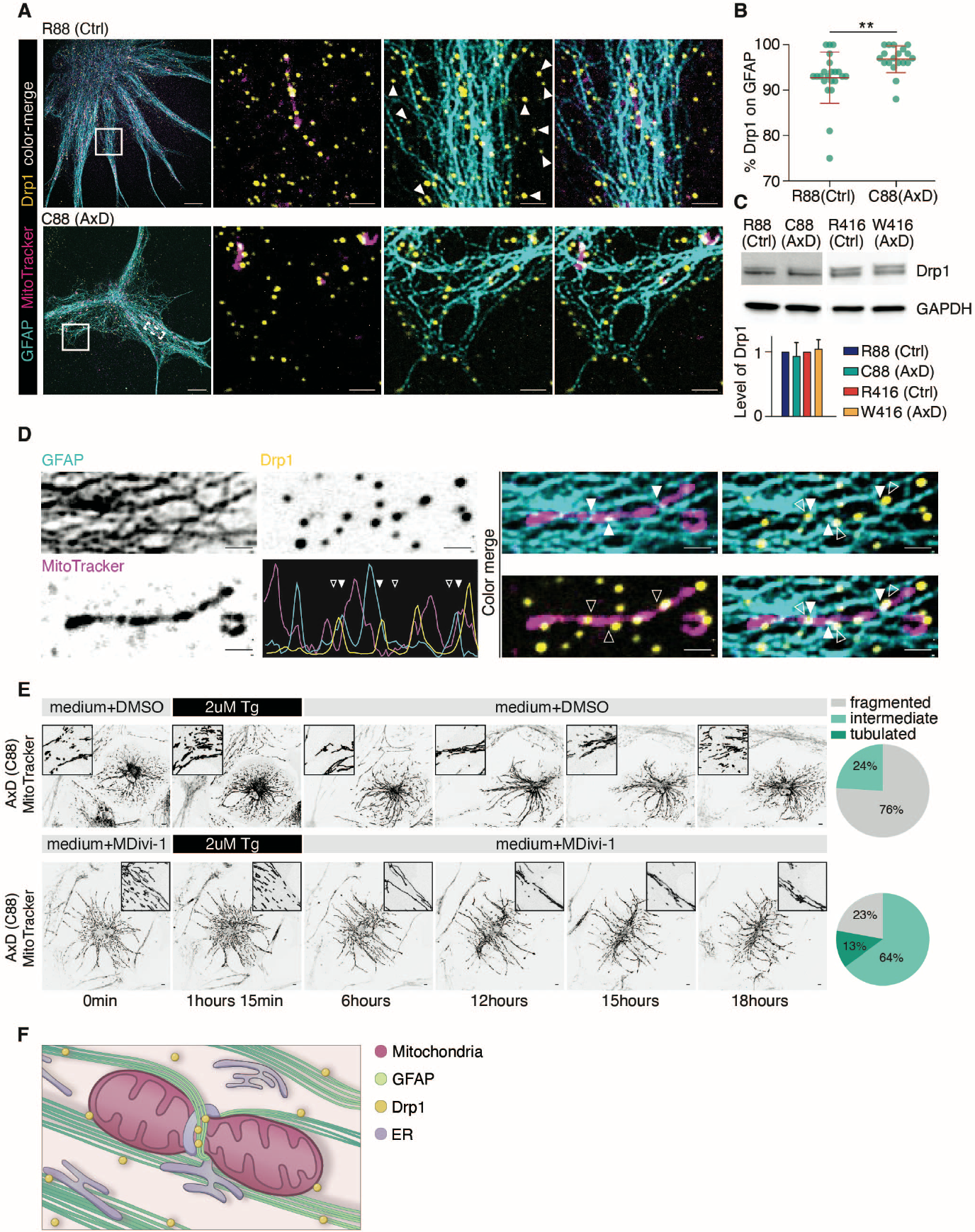
GFAP filaments serve as scaffolds for fission protein Drp1. (**A**) SIM images and high-magnification view of C88 AxD and isogenic control astrocytes immunostained for GFAP (Cell signaling technology, 34001) and Drp1, and stained with MitoTracker. White arrows indicate the Drp1 punctae that do not correlate with GFAP filaments. (**B**) Percentage of Drp1 puncta located on GFAP filaments (22 and 19 regions for control and AxD astrocytes, respectively). (**C**) Western blot and quantification showing the protein level of Drp1 in isogenic controls and AxD astrocytes (n=3 independent experiments). (**D**) High-magnification view and line scan analysis along mitochondria showing GFAP filaments and Drp1 puncta in mitochondrial constriction, originated from dashed line box in (C). Arrowheads in images and linescan highlight GFAP and Drp1 (white) positioned at mitochondria constriction sites (hollow). (**E**) Live-cell images and percentage of cells with indicated mitochondrial morphologies showing C88 AxD astrocytes mitochondrial change upon 2 μM thapsigargin. Cells are pre-incubated in medium with MDivi-1/DMSO, treated with thapsigargin and returned to medium with Mdivi-1/DMSO at indicated times (n=82 and 41 AxD cells for Mdivi-1 and DMSO incubation, respectively, 3 independent experiments). (**F**) Schematic representation showing the role of GFAP during mitochondria fission. **, p<0.005 by two-tailed unpaired *t*-test. Scale bar: confocal and SIM images, 10 μm, zoom-in SIM images, 2 μm.

If the GFAP-driven mitochondrial fission involves fission molecules, inhibition of Drp1 activity will reduce mitochondrial fission in the presence of an AxD-associated GFAP mutation. Indeed, pre-incubation of AxD astrocytes with the Drp1 inhibitor Mdivi-1^29,30^ rescued the fragmented mitochondrial phenotype, reducing the percentage of cells with fragmented mitochondria to 23%, closely resembling the isogenic control following thapsigargin washout (**Fig. 5E, movie. S6**). Additionally, it resulted in tubulated mitochondria in a subset (13%) of the AxD astrocytes (**Fig. 5E**).

In summary, our findings provide a previously unappreciated mechanism for mitochondrial fission where intermediate filaments drive mitochondrial constriction and serve as a scaffold for Drp1 (**Fig. 5F**). We provided two lines of evidence to support our model. First, GFAP filaments intersect mitochondria at fission sites under physiological condition throughout the entire constriction and fission process. Second, mitochondrial fission is blocked when GFAP is deleted under both physiological and stimulated conditions. While actin filaments have been suggested to facilitate mitochondria constriction before fission, several questions remain, such as what is the mechanism for selective binding of Drp1 to actin at fission sites and whether myosin provides the sole force for driving constriction^12^. Interestingly, we found that mitochondria fission is inhibited even upon Ca^2+^-induced stimulation in GFAP knockout astrocytes – this observation reinforces the indispensable role of GFAP despite the fact that there is acute recruitment of actin upon stimulation^11,31^.

Our observation is in agreement with the potential role of intermediate filaments as a force generation machinery suggested by the recent study of nuclear segmentation by vimentin^32^. The force generation could possibly occur through its interaction with other cytoskeletal component, like actin, to drive the constriction, as implied by the interpenetration between vimentin and actin filaments in the cell cortex^33,34^. Another possibility is the scaffolding mechanical support that intermediate filaments provide for organelles, such as the ER. It is known that ER tubules mark the sites of mitochondria division^5^, and vimentin serves as a scaffold for ER structure maintenance^35^. Indeed, we observed that the structure of ER was drastically altered in GFAP KO (**fig. S7**). Lastly, our model raises the question of whether GFAP’s role in mitochondria division is a general function shared by other types of intermediate filaments. It may provide a fundamental mechanism that explains/underlies neurodegeneration in Alexander disease resulting from GFAP mutations.

## Acknowledgments

We would like to thank Emma Feng for assistance with imaging.

## Funding

Singapore Ministry of Education, MOE2018-T2-2-103 (S-C.Z.)

Duke-NUS medical school, Duke-NUS-KPFA/2018/0022 (D.X.)

Singapore Ministry of Health, MOH-000207 and 000212 (S-C.Z.)

National Institute of Health, HD076892 (A.M.)

National Institute of Health, HD090256 (A.M.)

National Institute of Health, HD106197 (S-C.Z.)

Natural Science Foundation of China, 82301042 (D.X.)

Sichuan Science and Technology Program 2023ZYD0065 and 2023NSFSC0565 (D.X.)

## Author contributions

Conceptualization: SCZ, DX, LHK

Methodology: DX, LHK, YST

Investigation: DX, YST, LHK, XL, EA

Visualization: DX, YST, LHK, FY

Funding acquisition: SCZ, AM, DX

Supervision: SCZ

Writing-original draft: DX, SCZ, LHK, YST, FY

Writing-review and editing: DX, SCZ, LHK, ZJS, FY, AM

