## Supplementary materials for "The Alexander Disease Protein GFAP Drives Mitochondrial Fission"

**The PDF file includes:**

Materials and Methods

Figs. S1 to S7

**Other Supplementary Materials for this manuscript include the following:**

Movies S1 to S6

Materials and Methods

Reagents and antibodies

We used following live dye and chemicals: MitoTracker deep-red (Thermofisher), thapsigargin (Sigma), Mdivi-1 (Sigma), calcium indicator Fluo4-AM (Life technologies), ER-Tracker green (Thermofisher). We also used following antibodies: mouse anti-GFAP (monoclonal, Cell signaling technology, 34001), rat anti-GFAP 2.2B10 (monoclonal, Invitrogen, 13-0300), mouse anti-hGFAP STEM123 (monoclonal, Takarabio, Y40420), mouse anti-GFAP GA5 (monoclonal, Sigma-Aldrich, IF03L), goat anti-Sox9 (R&D, AF3075), rabbit anti-S100β (Abcam, ab52642), rabbit anti-Tomm20 (Thermofisher, MA532148), rabbit anti-Drp1 (Cell signaling technology, 5391S), mouse anti-GAPDH (Life technologies, MA515738), Hoechst 33342 (Life Technologies, H1399).

Plasmids and lentiviral packaging

The lentiviral vector (vector ID: VB210525-1200bqs) used to express GFAP-mNeon, pLV[Exp]-GFAP(short)>hGFAP[NM_002055.5](ns):3xGGGGS:mNeonGreen, was constructed and packaged by VectorBuilder. The C88 mutation was introduced into this plasmid using the In-Fusion® Snap Assembly kit (Takarabio) to generate a lentiviral vector for the expression of C88 mutant GFAP-mNeon. Lentiviral packaging plasmid psPAX2 (Addgene, #12260) and envelope plasmid pDM2.g (Addgene, #12259) were gifts from Didier Tronto. For lentiviral packaging, lentiviral vector, psPAX2 and pDM2.g were delivered into HEK293FT using Lipofectamine 3000 transfection reagent (Thermofisher), following the manufacturer’s instruction. Supernatants containing lentiviral particles were collected and filtered through a 0.45 μm membrane (Millipore, Bedford, MA) to remove cell debris, and pelleted by ultracentrifugation and resuspended in DMEM/F12 medium.

Cell lines and genomic editing

The CRISPR/CAS9 correction for C88 and W416 mutations were described previously^21^. H1 embryonic stem cell line was used to generate a GFAP knockout cell line using CRISPR/CAS9 technology. PAM site and sgRNA in the exon4 of human GFAP locus was identified with MIT CRISPR design tool (<http://crispr.mit.edu>). The sgRNA (AGAGATCCGCACGCAGTATG) was cloned into the pLentiCRISPR-V1 plasmid (Feng Zhang) and electroporated into H1 cells. Single clones were then screened for desired gene edition. Specifically, genomic DNA from each clone was extracted, DNA fragments flanking the sgRNA targeted site were amplified with PCR, Sanger-sequenced, and aligned with GFAP sequence from H1 cells. We identified multiple clones that harbour a single nucleotide knock-in mutation in both GFAP alleles, which results in frame-shift and an early stop codon 1bp downstream of the mutation site (still within the exon4). mRNAs containing premature termination codons (PTCs) are known to be degraded via nonsense-mediated mRNA decay (NMD), which results in the absence of protein product. This was validated by immunostaining and western blot in derived astrocytes.

Astrocyte differentiation, culture and transfection

All cell cultures were maintained in humidified incubators at 37°C with 5% CO_2_, and were tested negative for mycoplasma contamination. hPSCs were maintained as monolayer cultures on Matrigel in TeSR-E8 media (Stemcell technologies). The differentiation of astrocytes from hPSCs was described previously^22,21^. Neural differentiation was induced with dual SMAD inhibition (SB431542; DMH1)^36^. In brief, on day 0 of neural differentiation, the iPSC media was switched to neural differentiation medium (1/2 DMEM/F12+1/2 Neurobasal+0.5×N2 supplement+0.5×B27+1xNEAA +1 mM L-Glutamax), supplemented with the SMAD inhibitors SB431542 (2 μM) and DMH1 (2 μM). 14 days after the start of neural induction, neuroepithelia were lifted and propagated in neural medium (DMEM/F12+1× N2 Supplement+1× NEAAx+0.5x glutamax+2 μg/ml heparin) supplemented with FGF and EGF to generate astrocyte progenitors. Clusters of astrocyte progenitors were digested with TripLE (Gibco) and plated on 35-mm glass bottom culture dish (Cellvis), cover slip (Paul Marienfeld) or 6-well plate as single cells in 20,000 cells/cm^2^ density in medium supplemented with BMP4 (10 ng/ml) and CNTF (10 ng/ml) for terminal differentiation into mature astrocytes. For imaging, astrocyte progenitors were infected with lentivirus carrying WT-GFAP-mNeon or C88-GFAP-mNeon on day3 after the initiation of maturation.

Immunostaining

Astrocytes cultured on glass bottom dishes or coverslips were fixed in fresh 4% paraformaldehyde for 25min and washed with PBS for three times. Fixed cells were then permeabilized with 0.2% TritonX-100 for 10min and blocked with 10% donkey serum for 1 hour, followed by incubation with primary antibodies (diluted with 0.1% TritonX-100 and 5% donkey serum) overnight at 4°C. Primary antibodies were removed by washing with PBS for three times prior to the application of secondary antibodies (diluted in 5% donkey serum). After incubating for 30min at room temperature, cells were washed with PBS for three times. For imaging, cells on coverslips were mounted with Fluoromount-G mounting medium (Life Technologies, 00-4958-02), while cells cultured on glass bottom dishes were kept in PBS.

Transmission electron microscopy

Astrocytes growing on glass coverslip were fixed with 2.5% glutaraldehyde, 2.0% paraformaldehyde buffered in 0.1 M sodium phosphate buffer (PB). The samples were then post-fixed in osmium tetroxide and potassium ferrocyanide at the University of Wisconsin medical school electron microscopy facility, where the following steps including dehydration with ethanol series and acetone, infiltration with Durcupan ACM (Sigma-Aldrich) and acetone mixtures, embedding and polymerization in Durcupan ACM were performed. Samples were then cut into 100 nm sections on a Leica EM UC6 and imaged on a Philips CM120 transmission electron microscope equipped with an AMT BioSprint12 digital camera.

Confocal microscopy and SIM microscopy

Spinning disk confocal microscopy and super-resolution microscopy were performed on a Nikon Ti2-E inverted microscope with 4 laser lines (405 nm, 488 nm, 561 nm, 642 nm) equipped with a Yokogawa CSU-W1 T1 spinning disk confocal, a live super-resolution module (Live-SR; Gataca Systems) that is based on structured illumination (SIM) with optical reassignment and image processing^37^, and a Prime 95B sCMOS camera (Photometrics, pixel size 11 μmx11 μm). The method for super-resolution microscopy known as multifocal structured illumination microscopy (York et al., 2012) allows combining the doubling resolution with the physical optical sectioning of confocal microscopy, offering a two-time resolution improvement. The maximum resolution is 128 nm with a pixel size in super-resolution mode of 64 nm. Images were acquired using Nikon immersion objectives (Fluor 40x N.A. 1.3 oil, Apo 100x N.A. 1.49 oil). The microscope was controlled by Metamorph software (Molecular Devices). For all live imaging experiments, images were acquired at 37°C with 5% CO_2_ using an on-stage incubator and CO_2_ mixer (LCI), and cells were imaged in multi channels by sequential excitations with laser at 488 nm, 561 nm, or 642 nm through a quad-bandpass dichroic mirror (Semrock) and single band emitters (Semrock). Movies were acquired on a single z plane with speed in 3sec-5min/frame, and exposure times in 100–300ms range using either confocal or SIM microscopy. Images with whole cell z-stack were acquired with 0.5 μm step-size and maximum projection images of multiple z-stacks.

Ca^2+^ imaging

Astrocytes were cultured and matured on 35mm glass bottom dishes as described above. The cultures were maintained without medium change for at least 24h before thapsigargin experiments. Matured astrocytes were loaded with fluo4-AM diluted at a final concentration of 5 μM for 15 minutes before imaging. Single z plane images were recorded using confocal microscopy at 1frame every 3 seconds while thapsigargin diluted at final concentration of 2 μM was added during imaging. Fluorescence intensity was analyzed with a region of interest (30x30 pixels) after background subtraction, and fold change was quantified by (Ft_cell_-F0_cell_)/(F0_cell_-F0_background_).

Mitochondria dynamics synchronization

Astrocytes were cultured and matured on 35mm glass bottom dishes as described above, and loaded with 20 nM MitoTracker deep-red for 15 minutes before imaging. Single z plane images with multiple positions were recorded using confocal microscopy at 1 frame every 5 minutes. For thapsigargin induced mitochondrial fission experiments, images were acquired for 7 hours with recording a 20 minutes baseline before the addition of 2 μM thapsigargin. For mitochondrial dynamics synchronization, images were recorded for 22 hours with recording of 20 minutes baseline before addition of 2 μM thapsigargin. After thapsigargin incubation for 90 minutes, the medium was washed out and exchanged with fresh prewarmed medium during imaging; for rescuing the mitochondrial fragmentation in AxD astrocytes, culture medium was changed with medium containing 5 μM Mdivi-1.

Mitochondria fission event imaging

Astrocytes expressing WT/mutant-GFAP-mNeon were loaded with 20 nM MitoTracker deep-red for 15 minutes before imaging. Single z plane images were recorded using Live-SR SIM microscopy at 1frame every 3 seconds for 10 minutes. Dual-color images were acquired by sequential excitation of 491 nm and 642 nm lasers.

Image analysis

All images were analyzed using Fiji software. For the width of GFAP filaments and mitochondria, a line perpendicular to GFAP or mitochondria was drawn using Fiji “straight line” tool and “measure” was used for quantification. The level of colocalization of mitochondria over GFAP was analyzed by calculating the mander’s coefficient using “JACOP” plugin. To confirm the overlap is not due to chance, mitochondrial images were rotated 90° counter-clockwise, and the mander’s coefficient was quantified. The percentage of mitochondria correlated with GFAP was quantified by manually counting the number of mitochondria attached to GFAP. For the morphology of mitochondria, we scored mitochondria as in a previous study^23,24^. To measure individual mitochondrial length, maximum intensity projections of z-series images with 0.5 μm increments for MitoTracker were generated. A line along mitochondria was drawn using the Fiji “freehand line” tool, and the “measure” function was used for quantifying length. The number and length of all the mitochondria in an individual cell, except for the center region where mitochondria stacked together, were quantified by two independent observers blind to the experiments. For the dynamics of mitochondrial morphology after thapsigargin synchronization, three time points with mitochondrial fusion after thapsigargin washout, return back to intermediate status, and return back to fragmented status (especially AxD astrocytes) were recorded and plotted. The corresponding mitochondrial morphology and individual mitochondrial length were analyzed at five timepoints as indicated. To measure the percentage of Drp1 puncta associate with GFAP, the number of Drp1 puncta on 200x200 pixel regions from single z-plane SIM images were used for quantification. The percentage was quantified by number _Drp1 puncta on GFAP_/ number _total Drp1 puncta_. For identifying GFAP filaments during mitochondrial constriction and fission by linescan, a line was drawn through mitochondria perpendicular to the fission site, and the fluorescence intensity of GFAP and mitochondria was measured along the length of line for each time point and plotted.

Western blotting

Astrocytes were washed with cold PBS, scratched and lysed on ice using RIPA lysis buffer (Thermo Fisher) together with protease inhibitor, phosphatase inhibitor, PMSF and Dithiothreitol. Total protein concentration was measured by BCA protein assay. 4X Laemmli sample buffer was added into the protein lysate and boiled at 100℃ for 5 minutes. Protein samples were loaded on 4%-20% Mini-Protein SFX precast gel (Bio-rad), transferred to polyvinylidene difluoride membranes, blocked with 5% non-fat dry milk TBST, and incubated with primary antibodies overnight at 4°C. Signals were visualized using horseradish peroxidase-conjugated secondary antibodies, ECL system and captured with ChemiDoc system.

Statistics

Statistical analysis was evaluated by paired/unpaired two-tailed Student’s *t*-test as indicated in figure legends. Average data is shown as means ±S.D. All statistical analysis was performed using GraphPad Prism software.

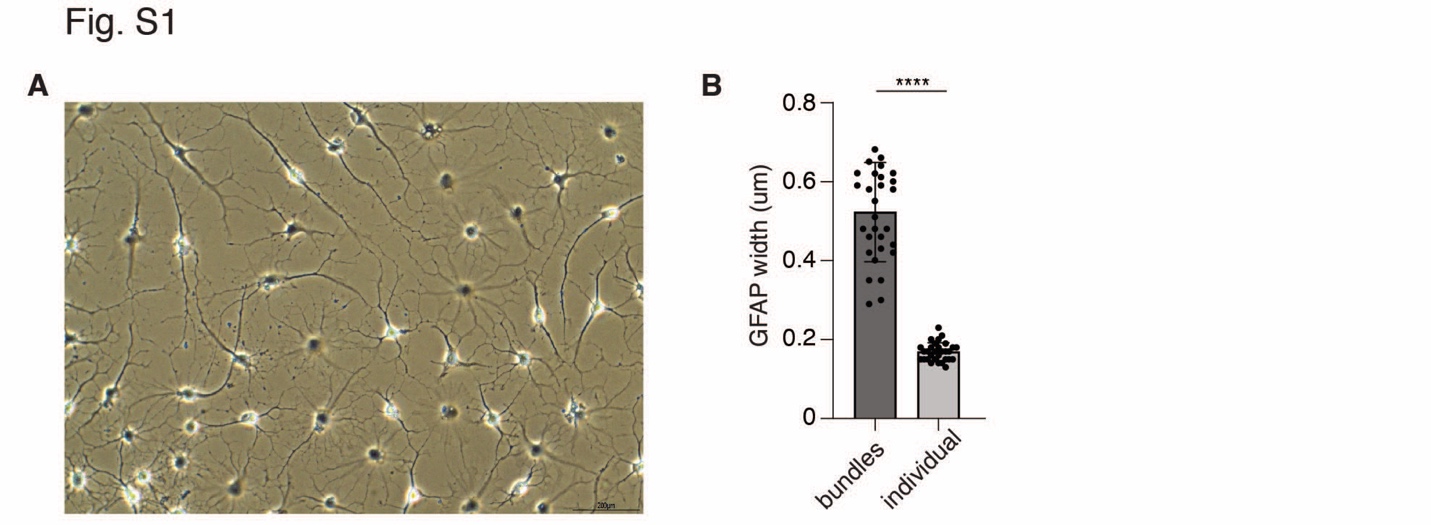

Fig. S1. Characteristics of astrocyte morphology and GFAP filaments. (A) Representative bright field image of astrocytes matured for 14 days. (B) Quantification of the width of GFAP bundles and individual filaments under SIM microscopy (n=33 filament bundles and 30 individual filaments, 4 independent experiments). ****, p ≤0.0001 by two-tailed unpaired *t*-test. Scale bar, 200 μm.

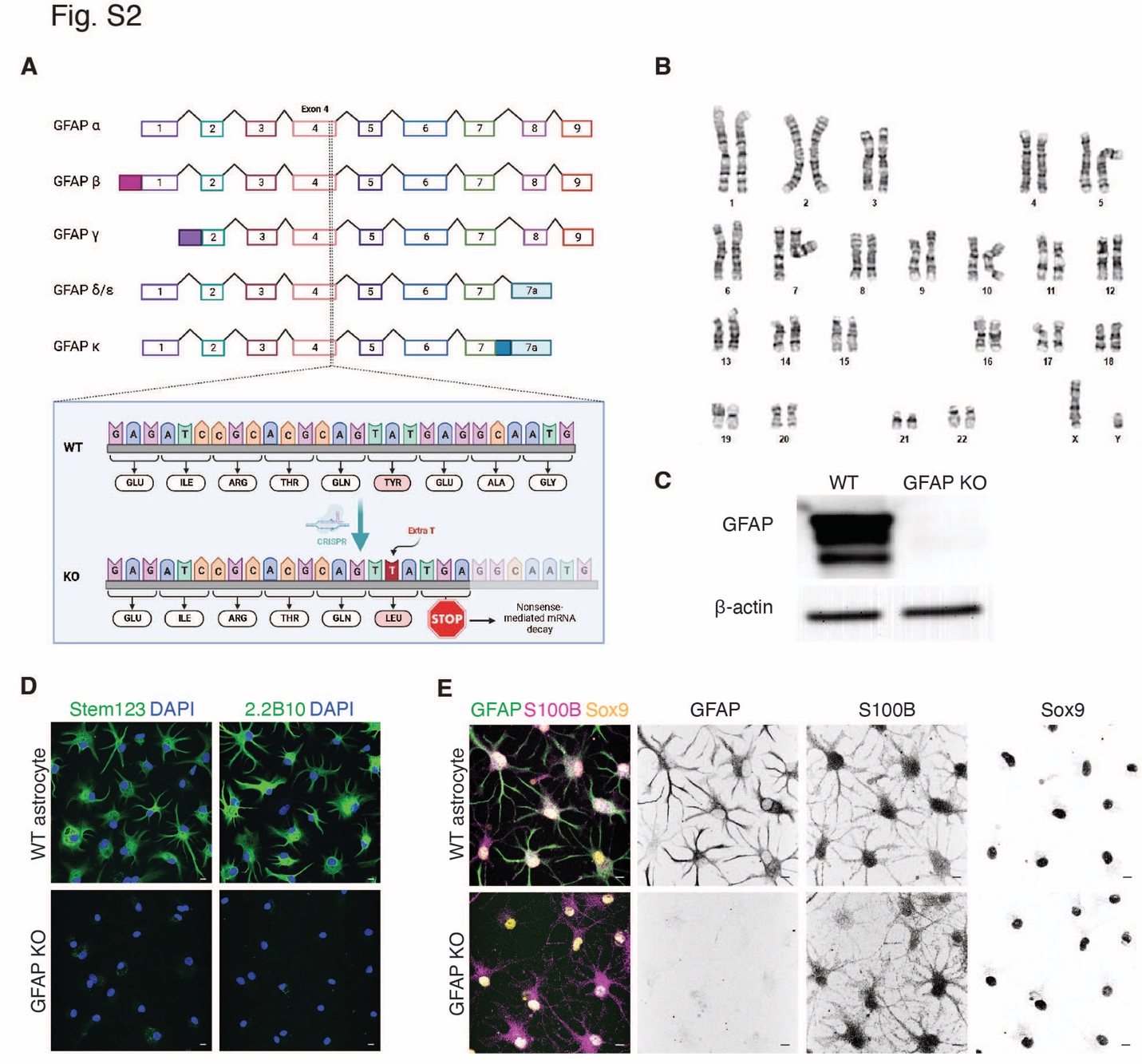

Fig. S2. Generation and validation of GFAP knockout astrocytes from hPSCs. (A) Schematics showing the generation of GFAP knockout cells from hPSCs. (B) Representative karyotyping results from a GFAP knockout line. (C) Western blotting showing the expression level of GFAP in GFAP KO and parental astrocytes immunostained with mouse anti-GFAP GA5 (Sigma-Aldrich, IF03L), 3 independent experiments. (D) Confocal images showing GFAP knockout and parental astrocytes immunostained with two GFAP antibodies mouse anti-hGFAP STEM123 (Takarabio, Y40420) and rat anti-GFAP 2.2B10 (Invitrogen, 13-0300), 3 independent experiments. (E) Confocal images showing the GFAP knockout and parental astrocytes immunostained for common astrocyte markers GFAP (rabbit anti-GFAP, DAKO, Z0334), S100β, and Sox9 (3 independent experiments). Scale bar, 10 μm.

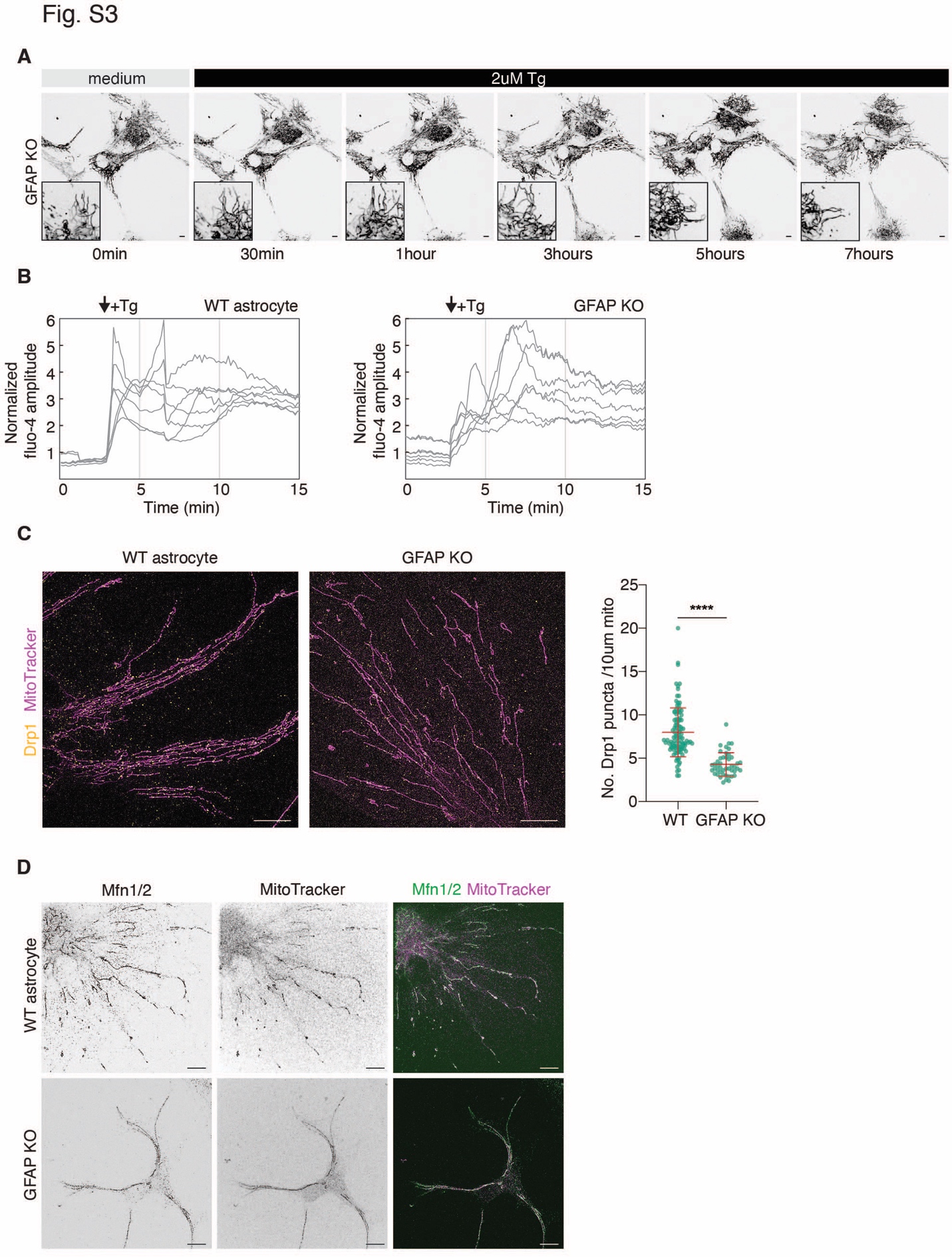

Fig. S3. Mitochondrial dynamics, calcium signals and Drp1 staining in GFAP knockout astrocytes. (A) Another example of live-cell images and magnified images of GFAP knockout astrocytes stained with MitoTracker when treated with 2 μM thapsigargin. (B) Intensity plots showing cytosolic calcium changes in wildtype and GFAP knockout astrocytes upon 2 μM thapsigargin. Astrocytes are stained with calcium dye Fluo-4 (3 independent experiments). (C) SIM images of wildtype and GFAP knockout astrocytes stained with MitoTracker and antibody against Drp1, and quantification of Drp1 puncta per mitochondrial length (n=107 and 46 mitochondria from wildtype GFAP knockout astrocytes, respectively, 3 independent experiments). (D) SIM images of wildtype and GFAP knockout astrocytes stained with MitoTracker and antibody against Mfn1/2 (n=13 wildtype and 15 GFAP knockout astrocytes, 3 independent experiments). ****, p ≤0.0001 by two-tailed unpaired *t*-test. Scale bar, 10 μm.

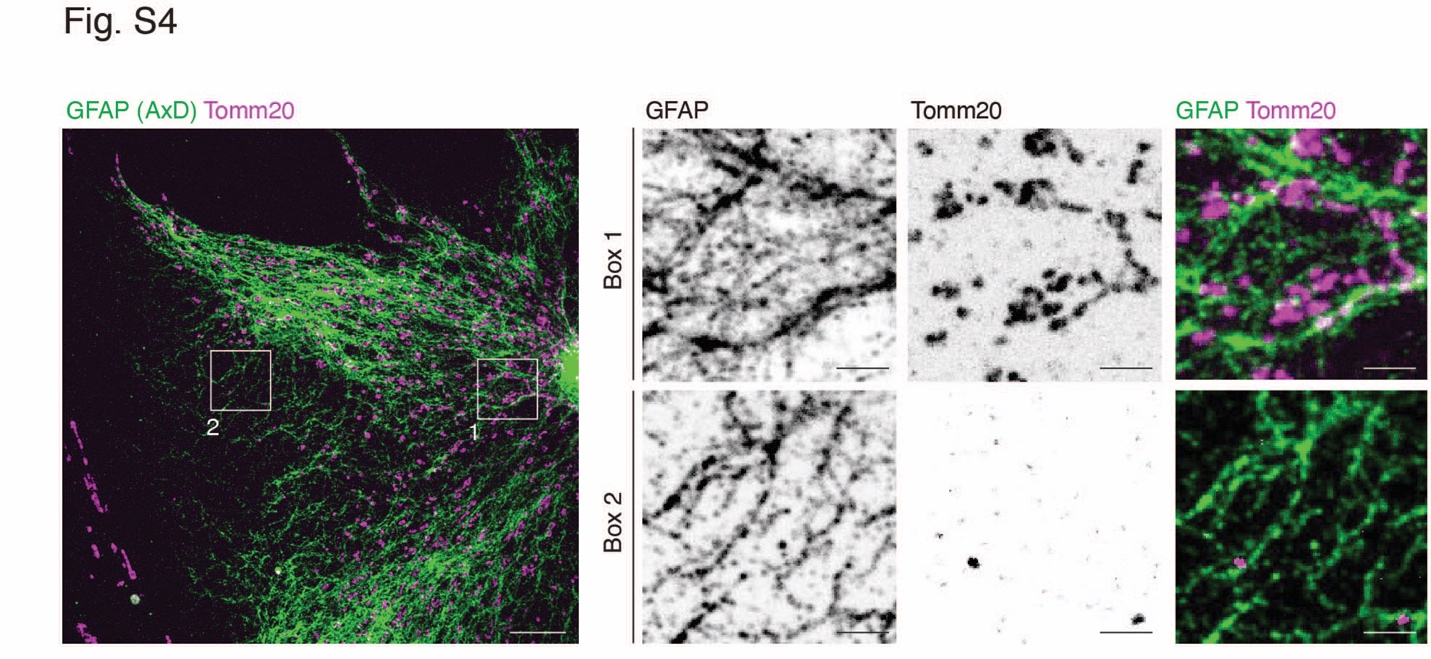

Fig. S4. Mitochondrial distribution and their spatial relation with GFAP in AxD astrocytes. Representative SIM image of C88 astrocytes co-immunostained with antibodies against GFAP (green) and Tomm20 (magenta). Two insets are amplified to show the relationship between GFAP filaments and mitochondria. Scale bar: SIM images, 10 μm; zoom-in SIM images, 2 μm.

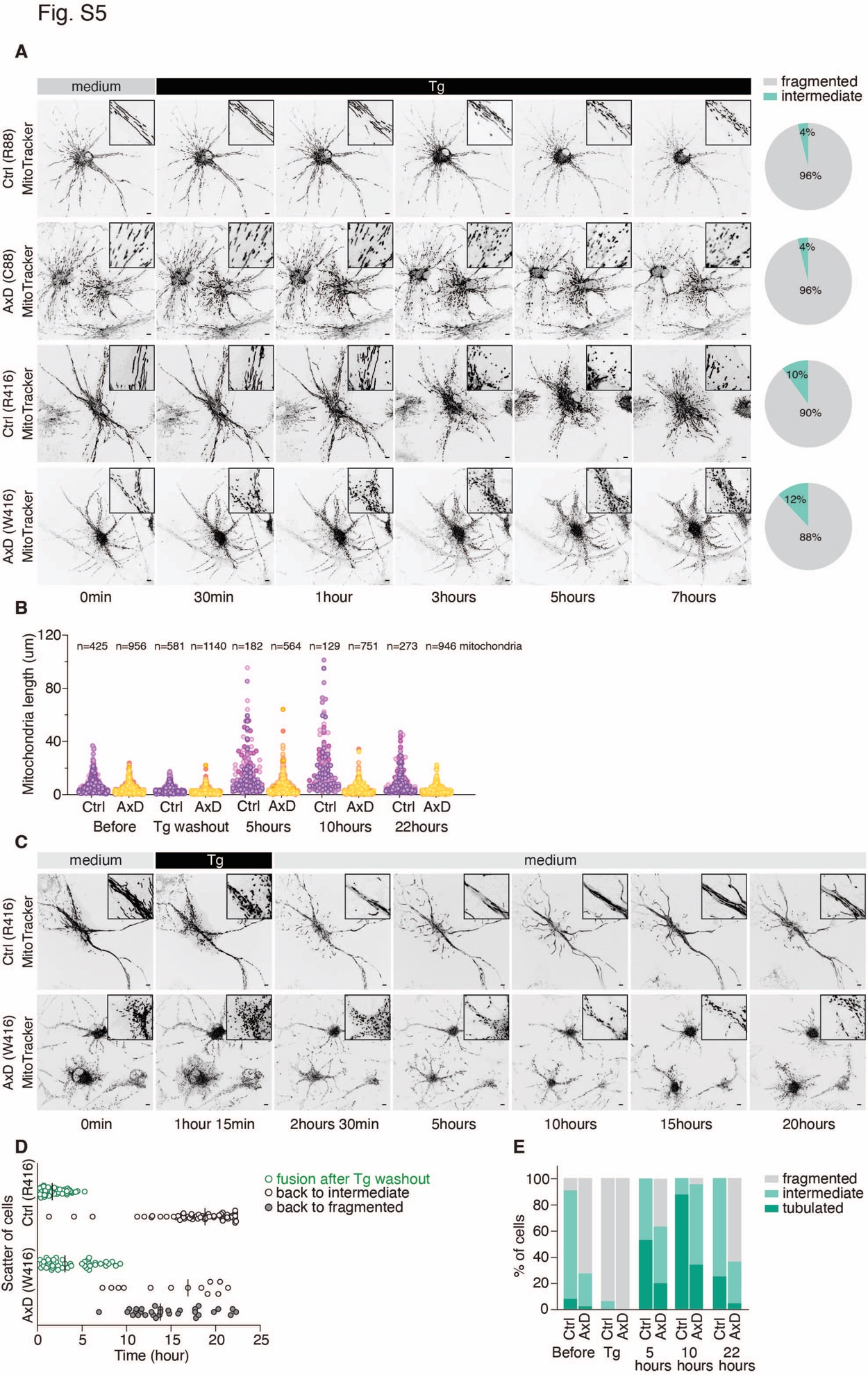

**Fig. S5. Morphological changes of mitochondrial in AxD and isogenic control astrocytes upon thapsigargin.** (**A**) Live-cell images of mitochondria (indicated by MitoTracker) in AxD and isogenic control astrocytes that are treated with 2 μM thapsigargin at indicated times. The percentage of cells with indicated mitochondrial morphologies are shown in the right panel (94 R88 cells, 152 C88 cells, 101 R416 cells and 97 W416 cells from 3 independent experiments). (**B**) Plot of individual mitochondrial length for R88 and C88 astrocytes at indicated time (numbers of mitochondria are listed in figure, 4 cells for R88 and C88 cells, respectively). (**C**) Live-cell images showing the dynamics of mitochondrial fission-fusion in R416 and W416 astrocytes. Astrocytes stained with MitoTraker are synchronized by 2 μM thapsigargin and changed back to normal medium as indicated. Images are taken at 1 frame every 5 minutes (n=64 R416 cells and 44 W416 cells, 3 independent experiments). (**D**) Scatter plot shows the time point at which mitochondria in R416 and W416 astrocytes start fusion after thapsigargin washout (green), return back to intermediate mitochondria status (black circle) or further go back to fragmented status (grey). Each dot represents one cell, with average time shown as black lines. (**E**) Percentage of R416 and W416 astrocytes in tubulated (green), intermediate (light green) and fragmented (grey) status at indicated times. Scale bar, 10 μm.

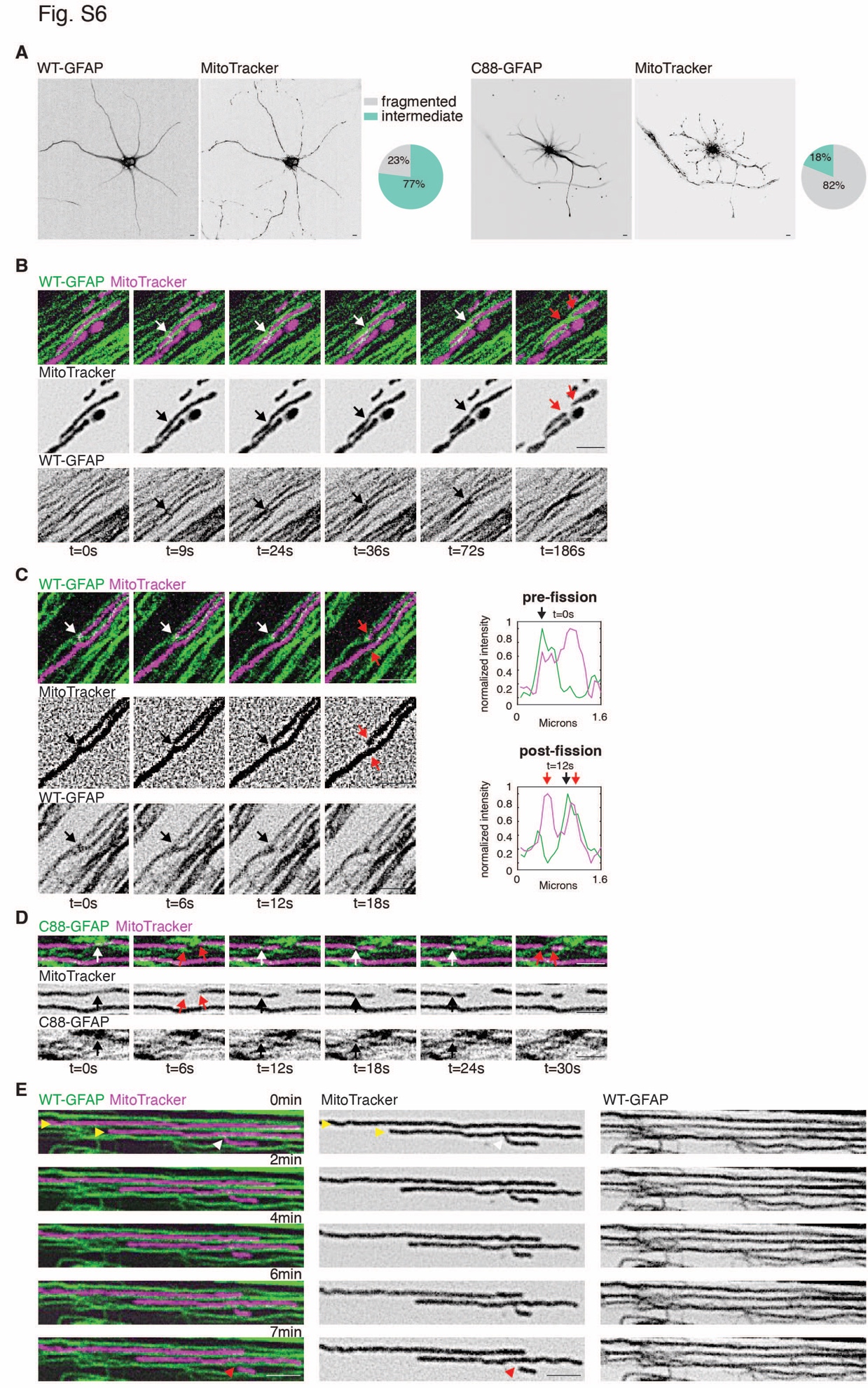

**Fig. S6.** **Dynamics of GFAP filaments in mitochondrial fission by live super-resolution imaging.** (**A**) Representative confocal images and percentage of cells with indicated mitochondrial morphologies in isogenic control astrocytes expressing WT or C88-GFAP-mNeon, and stained with MitoTracker. (70 and 66 cells for WT- and C88-GFAP- mNeon expressing, respectively, 3 independent experiments). (**B**) Example of magnified SIM live-imaging showing WT-GFAP filaments crossing, sliding and encircling mitochondria (white arrow) prior to mitochondrial fission (red arrow). (**C**) Left, another example of SIM live-imaging showing WT-GFAP filaments crossing the site of mitochondrial division (white arrow) before fission (red arrow); right, corresponding line scans analyze the fluorescence intensity of mitochondria and GFAP pre- and post-fission. (**D**) Magnified SIM live-imaging showing C88-GFAP crossing the site of mitochondrial division (white arrows) before two fission events (red arrows). (**E**) Magnified SIM live-imaging showing that mitochondria aligned along GFAP filaments (yellow arrowhead) do not undergo fission. In contrast, mitochondria undergo fission (red arrowhead) at a position crossed by GFAP (white arrowhead). Scale bar: confocal images, 10 μm; zoom-in SIM images, 2 μm.

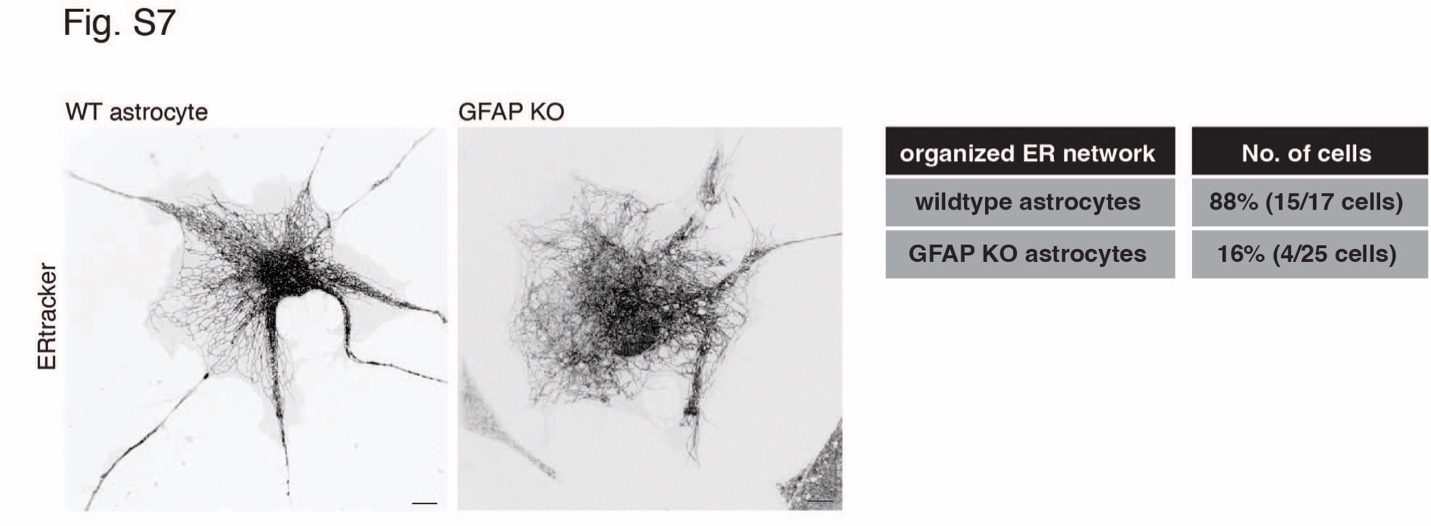

Fig. S7. Disruption of organized ER network in GFAP knockout astrocytes. Representative confocal images and percentage of cells with organized ER network from H1 wildtype and GFAP knockout astrocytes stained with 1μM ER-Tracker (n=17 wildtype astrocytes and 25 GFAP knockout astrocytes). Scale bar: 10 μm.

Movie S1. SIM video of wildtype and GFAP knockout astrocyte stained with MitoTracker showing mitochondrial changes in the presence of 2μM thapsigargin. The video was acquired at 5 minutes interval and played at 30 frame per second (9000x real time). Scale bar: 10 μm.

**Movie S2.** SIM video of C88, W416 AxD astrocytes and their isogenic control R88 and R416 astrocytes stained with MitoTracker showing mitochondria fragmentation upon 2 μM thapsigargin treatment. The video was acquired at 5minutes interval and played at 30 frame per second (9000x real time). Scale bar: 10 μm.

Movie S3. SIM video of C88 AxD and isogenic control R88 astrocytes stained with MitoTracker showing the dynamics of mitochondrial morphology. Astrocytes were synchronized by 2 μM thapsigargin and changed back to normal medium as indicated. The video was acquired at 5 minutes interval and played at 30 frame per second (9000x real time). Scale bar: 10 μm.

Movie S4. High magnification of two-color SIM videos of astrocyte expressing WT-GFAP-mNeon and stained with MitoTracker showing the participation of GFAP filament during mitochondrial fission. Three representative fields of view are shown. Videos were acquired at 3 seconds interval and played at 20 frame per second (60x real time). Scale bar: 1 μm.

Movie S5. High magnification of two-color SIM video of astrocyte expressing mutant C88-GFAP-mNeon and stained with MitoTracker showing the participation of GFAP filament during mitochondrial fission. Two representative fields of view are shown. Videos were acquired at 3 seconds interval and played at 10 frame per second (30x real time). Scale bar: 1 μm.

Movie S6. SIM video of C88 AxD astrocytes stained with MitoTracker showing the dynamics of mitochondrial morphology upon 2 μM thapsigargin. Astrocytes are pre-incubated in medium with 5 μM MDivi-1, treated with thapsigargin and return back to medium with MDivi-1 at indicated times. The video was acquired at 5 minutes interval and played at 30 frame per second (9000x real time). Scale bar: 10 μm.
